# A novel newborn screening modality: Non-targeted proteome analysis using low-cost iron powders

**DOI:** 10.1101/2024.12.25.630353

**Authors:** Daisuke Nakajima, Masaki Ishikawa, Ryo Konno, Hideo Sasai, Osamu Ohara, Yusuke Kawashima

**Affiliations:** Department of Applied Genomics, Kazusa DNA Research Institute, 2-5-23 Kazusa Kamatari, Kisarazu, Chiba 292-0818, Japan

**Keywords:** dried blood spots, inherited diseases, newborn screening, non-targeted proteome analysis

## Abstract

In this study, we developed a simple protein extraction method for dried blood spots (DBS) that potentially meets the throughput required for newborn screening (NBS) and optimizes non-targeted proteomic analysis in combination with liquid chromatography coupled mass spectrometry in the data-independent-acquisition mode (DIA-LC-MS/MS). The developed pipeline, termed **<u>N</u>**on-targeted **<u>A</u>**nalysis of **<u>N</u>**on-specifically **<u>D</u>**BS-**<u>A</u>**bsorbed proteins (NANDA), successfully addressed the following three challenges: (1) processing of 96 3.2-mm DBS punches in parallel using low-cost iron powders with a robotic system, (2) identifying more than 5,000 proteins using DIA-LC-MS/MS, and (3) improving DIA-LC-MS/MS throughput to 40 samples/day with minimal compromise in protein coverage depth. The results imply that this pipeline can open new venues for conducting NBS using non-targeted quantitative proteome profiling, which has been a currently missing modality in NBS.

## Introduction

Blood components are an important source of biomarkers for disease detection because they are strictly kept within a normal range by homeostasis in the healthy state. In fact, many diagnostic tools using fresh whole blood, serum, or plasma have been widely utilized. As a modified approach, dried blood spots (DBS) have been used as an alternative to fresh blood because of the simplicity of shipping and storage (McDade *et al*, 2007; Li & Tse, 2010; Keevil, 2011; Deglon *et al*, 2012). For example, DBS has served as a versatile format for newborn screening (NBS) to prevent disease onset and/or reduce symptoms by early intervention. The DBS approach is well-established and has been proven effective in conventional NBS based on the measurements of specific metabolites or enzymatic activities (Saudubray & Garcia-Cazorla, 2018). As the number of treatable diseases increases with the emergence of new treatments, new NBS modalities have been introduced. Examples include the copy number measurement of the SMN1 gene and T/B-cell excision circle DNA for spinal muscular atrophy (Nishio *et al*, 2023) and severe combined immunodeficiency (Currier & Puck, 2021), respectively. However, the increase in screening items inevitably leads to increased screening costs and labor, presenting a realistic and serious concern. To address this issue, we are actively seeking an appropriate “one-size-fits-all” screening modality. To this end, we propose measuring biomolecular profiles in a non-targeted manner, followed by data filtering, as a solution. Similarly, non-targeted genome sequencing analysis has gained attention as a new NBS modality. This non-targeted approach could enable us to conduct NBS for multiple diseases using a “one-size-fits-all” platform. However, although genome sequencing may serve as a versatile screening approach for various inherited diseases, various debates regarding its suitability for NBS, including ethical, economic, and social issues, are currently underway. Additionally, a concern arises that genomic information, which pertains to genotype, does not fully convey the molecular phenotype of neonates, on which current NBS is based before clinical symptoms appear. In this context, we propose non-targeted quantitative protein profiling as a novel NBS modality, offering an alternative to genome sequencing.

In a proof-of-concept study, we recently demonstrated that a non-targeted proteomic approach using data-independent acquisition liquid chromatography-tandem mass spectrometry (DIA LC-MS/MS) on DBS could offer complementary insights into personal genome sequences for diagnosing inborn errors of immunity (Shibata *et al*, 2024). However, the pipeline used by Shibata *et al*. was labor-intensive, and the overall throughput was not high enough for NBS, implying that a modified pipeline for the non-targeted proteome analysis of DBS must be developed to move beyond the proof-of-concept stage. In this study, we thus set the following goals to be achieved by a new pipeline: 1) capability of the parallel processing of 96 DBS punches using automation at low cost, 2) deeper DBS proteome coverage than that in our previous method, and 3) improvement of overall throughput to 10, 000 samples/year/LC-MS instrument. By introducing several new methodologies, we successfully constructed a new pipeline that robotically extracts proteins from 96 individual DBS punches in parallel with a protein coverage of 5,000 at a throughput of 40 DBSs/day. We believe the developed DBS- manipulation pipeline paves the way for introducing non-targeted protein profiling as a new modality in NBS.

## Results and Discussion

### Proof-of-concept experiment of the non-targeted proteome analysis of filter paper-retained proteins

In our previous study, we removed abundant soluble serum/plasma proteins from DBS extracts using salt-induced precipitation (Nakajima *et al*, 2020). However, we considered further improvements in protein coverage depth and throughput mandatory for implementing non-targeted proteome analysis in real-world NBS. In the trial and error processes to this end, we noticed that considerable amounts of proteins remained on DBS even after extensive washing with buffer solution. This finding suggested that we could use the filter paper used for DBS as a solid support that absorbs proteins (probably with low solubility) non-specifically. If this is the case, simple washing of DBS with an appropriate buffer might enable us to remove a large amount of soluble serum/plasma proteins without losing low-solubility proteins on the filter paper. To pursue this possibility, we conducted a proof-of-concept experiment as described below.

In this context, we devised a **<u>N</u>**on-targeted **<u>A</u>**nalysis of **<u>N</u>**on-specifically **<u>D</u>**BS-**<u>A</u>**bsorbed proteins (NANDA), as shown in Fig 1. In NANDA, DBS was first suspended in tris buffered saline with Tween 20 (TBST) until it became a suspended paper pulp and then centrifuged to precipitate the paper pulp. Next, the precipitate was washed once with TBST and twice with 50 mM Tris-HCl pH 8.0 to separate the filter paper debris from which soluble proteins such as albumin had been removed. Since a considerable amount of proteins was kept bound on the filter paper pulp even after extensive washing with TBST, we directly added protease to this pulp suspended in 50 mM Tris-HCl pH 8.0 for digestion and finally yielded approximately 2–5 µg of digested peptides in the suspension. The DIA-LC-MS/MS data of the peptide mixtures thus obtained indicated that NANDA allowed us to identify approximately 1.8-fold (3,541/1,925) more proteins (the number of protein groups stably detected in four replicates by each method) than the sodium carbonate precipitation (SCP) method reported in 2020 (Nakajima *et al*., 2020) (Fig 2A). Over 93% (1,809 out of 1,925) of the proteins identified by the SCP method were also detected using NANDA (Fig 2B).

**Figure 1.**
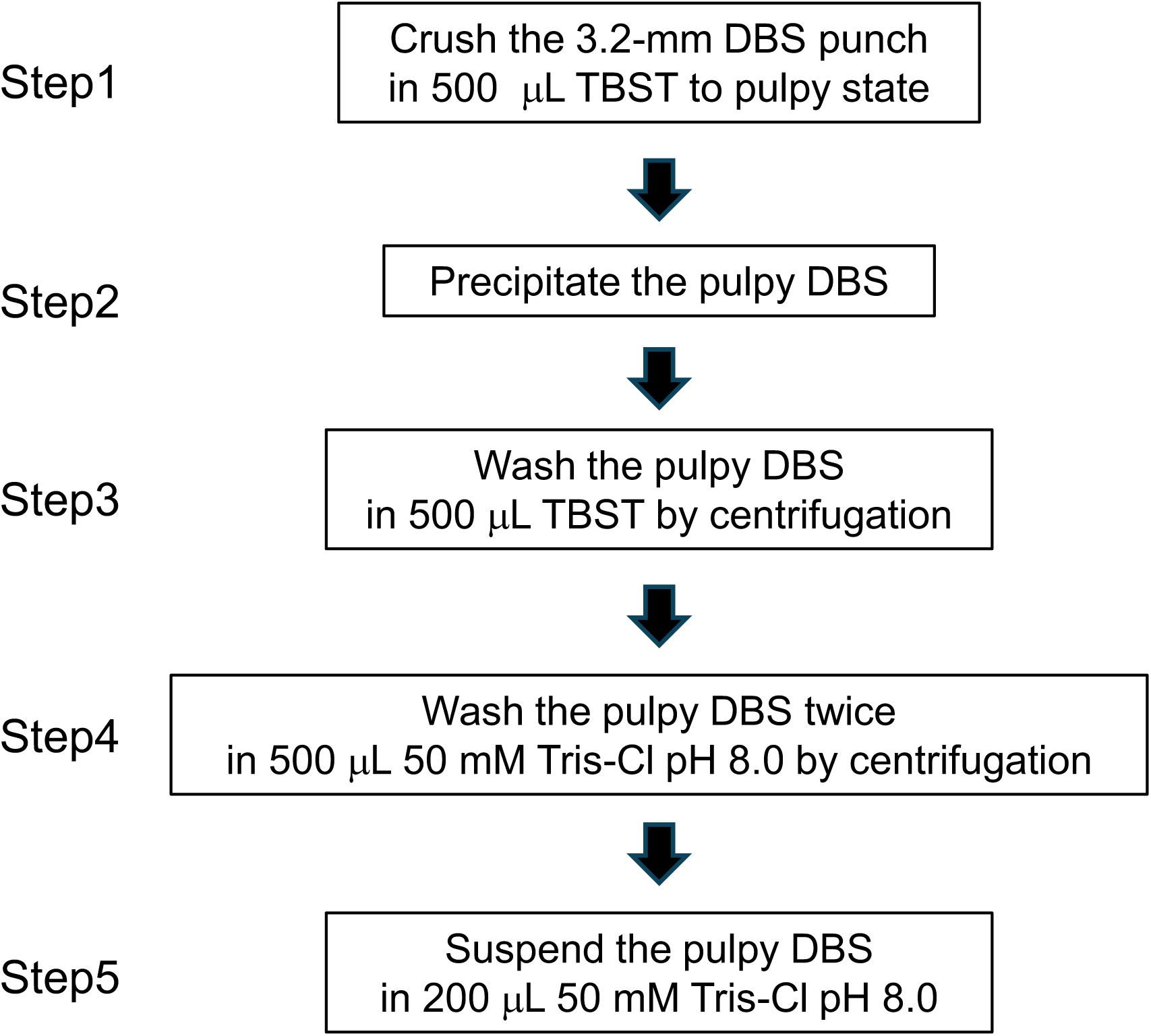
Flowchart of the preparation of non-targeted analysis for non-specifically dried blood spot-absorbed proteins.

**Figure 2.**
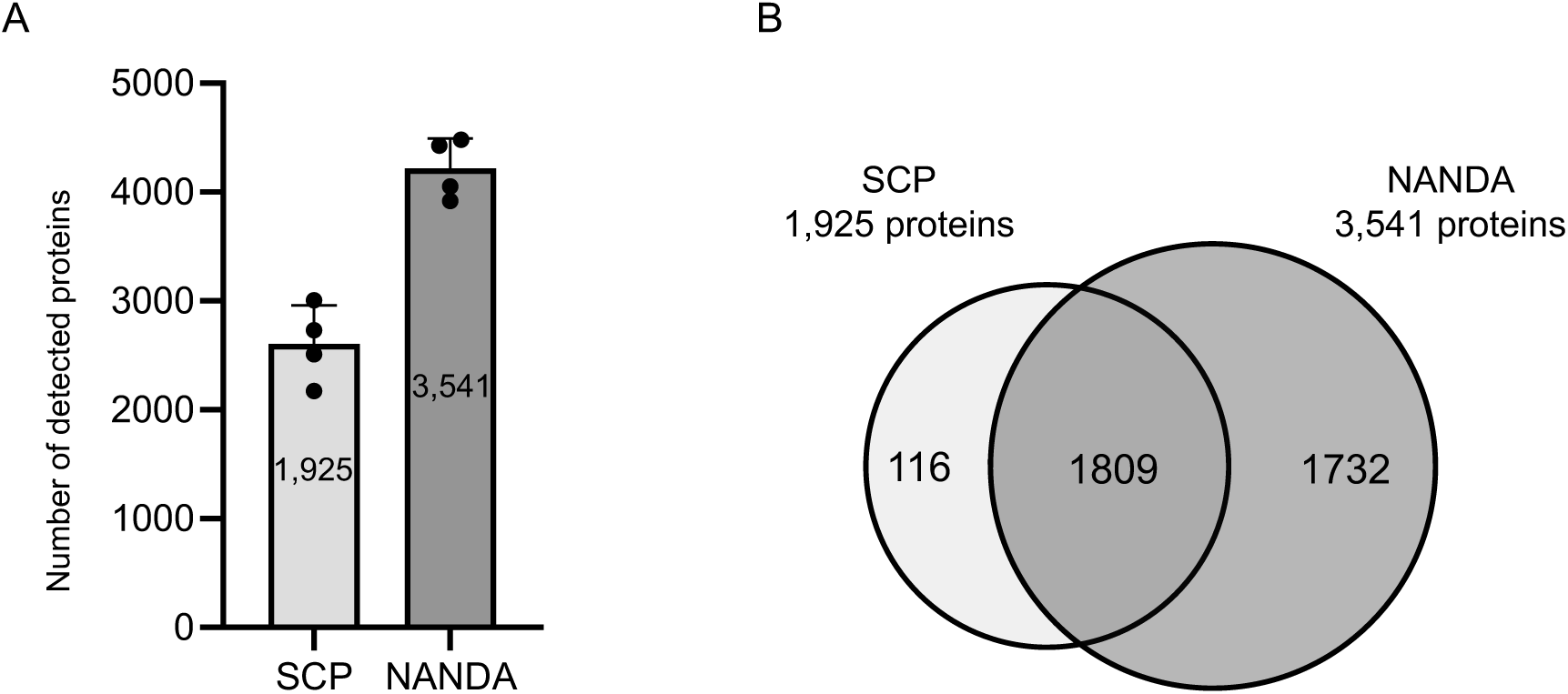
Comparison of the number of proteins detected using sodium carbonate precipitation and non-specifically dried blood spot-absorbed proteins. A Bar graph showing the number of proteins detected using sodium carbonate precipitation (SCP) and non-specifically dried blood spot-absorbed proteins (NANDA) (*n* = 4). The number in the bar graph shows the number of proteins commonly detected in four replicates. B Venn diagram comparing the proteins detected by the two preparations. The number in the Venn diagram indicates the number of proteins commonly detected in four replicates.

The data described above demonstrated that NANDA was simpler and less labor-intensive but provided deeper protein coverage than did the SCP method. In this regard, it should be noted that analysis of non-specifically bound proteins on DBS is not a new concept in literature: Molloy *et al*. reported that a simple DBS washing step using a sophisticated volumetric absorptive microsampling device (VAMS) made it possible to robustly quantify up to 1,600 proteins from single-shot shotgun LC-MS/MS analysis (Molloy *et al*, 2022). The methodological differences between NANDA and their report include differences in solid support (conventional filter paper vs VAMS support) and washing conditions. As for the solid support, we confirmed that NANDA provided the same results using conventional filter papers used for NBS supplied by three different suppliers (Fig EV1), although we did not check the solid support we used in VAMS. As for the washing conditions, DBS was subjected to more extensive washing in NANDA as a suspended paper pulp than in the method by Molly *et al*. We assume that the complete removal of contaminating abundant plasma proteins must contribute to deepening the identified protein coverage compared with that in the method by Molly *et al*. Together with optimizing the precursor MS range and isolation window width (Fig EV2), the number of identified protein groups exceeded 5,000 using NANDA coupled with the DIA-LC-MS/MS system.

After confirming the concept of NANDA, we next sought to optimize and develop a robotic system for NANDA to improve overall throughput. The characteristics of protein groups detected using NANDA are described in detail in the subsequent sections.

### Development and validation of automated NANDA platform

#### Centrifugation-free washing of filter paper pulp using low-cost iron powders

To improve NANDA throughput, we attempted to develop a centrifugation-free system. Because the inclusion of high-speed centrifugation in NANDA made building a high-throughput sample-processing platform difficult, we aimed to develop an alternative to wash dispersed paper pulp without centrifugation steps. After much trial and error, we finally arrived at a method that enabled manipulating suspended paper pulp using magnetism instead of centrifugation. In this system, we used iron powder as a cost-effective magnetic carrier to tie the suspended paper pulp into a bundle. The video showing this process is available in Expanded View (Video EV). Next, we implemented the iron powder-assisted NANDA (auto-NANDA) on a robotic processing device for magnetic beads (Maelstrom 8-Autostage or Maelstrom 9610; TANBead, Taoyuan City, Taiwan). Fig 3A shows the auto-NANDA scheme. By adding iron powder during DBS suspension in the initial TBST, a mixture of iron powder and dispersed paper pulp was produced automatically by the rotational motion of the spin tip. By inserting a magnetic rod into the spin tip, this complex was magnetically bound to the spin tip, allowing for easy and almost complete isolation. The iron powder and paper pulp composite is bound more compactly to the spin tip than to the centrifugation-assisted collection of suspended paper pulp, resulting in less carryover of washing solution retained by the paper pulp. This certainly improved the washing efficiency (Fig 3B). The composite was then transferred to a new well and washed once with TBST and twice with 50 mM Tris-HCl pH 8.0 using the spin tip’s rotation and magnetic manipulation. Finally, the composites were robotically transferred to a trypsin-digestible solution. Once the reagents were set in a 96-well plate, the entire process took approximately 1 h, with Maelstrom 8-Autostage processing for 8 samples and Maelstrom 9610 processing for 96 samples simultaneously.

**Figure 3.**
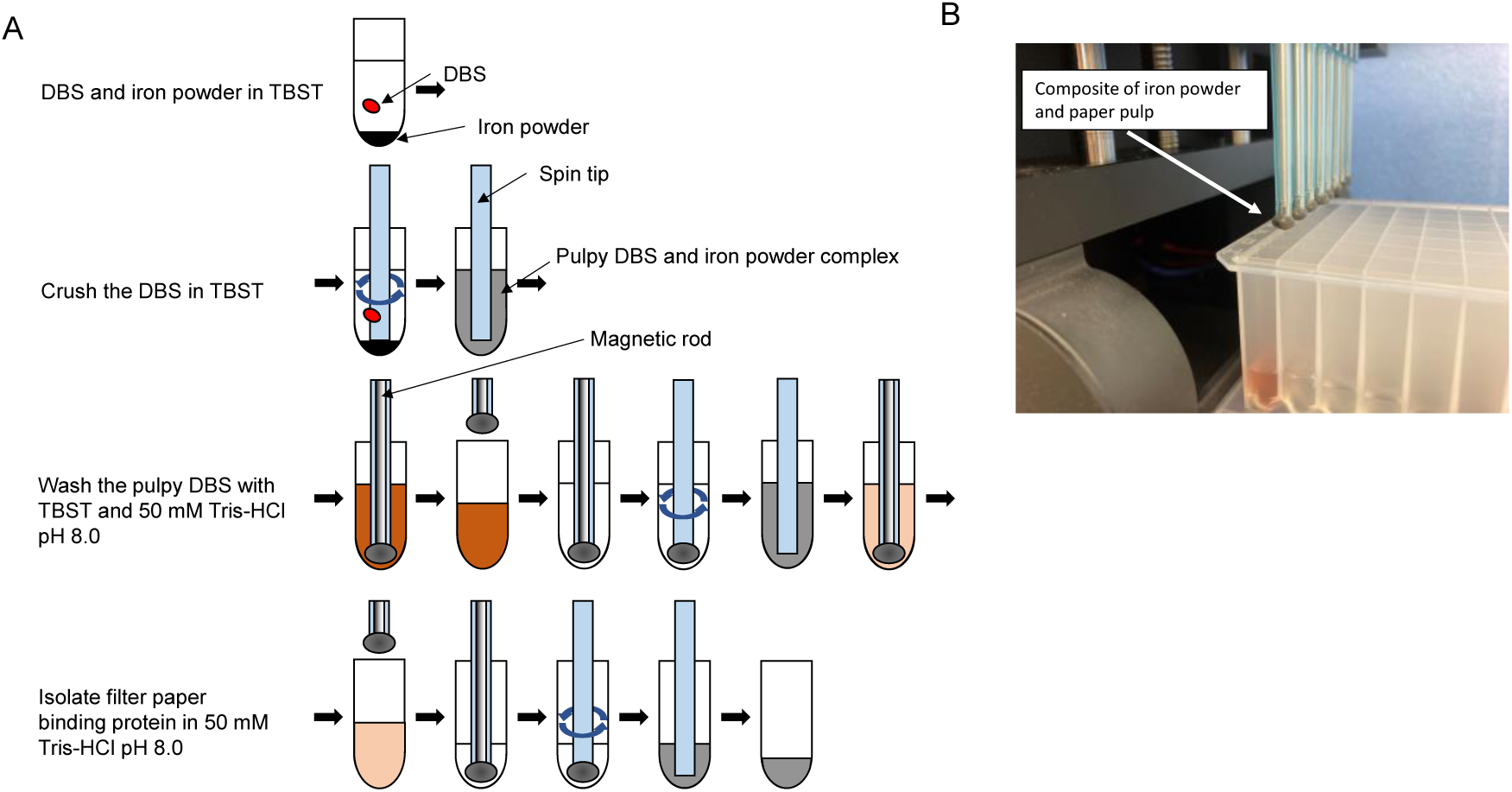
Preparation of automated non-specifically dried blood spot-absorbed proteins. A Schematic illustration of the preparation of automated non-specifically dried blood spot-absorbed proteins using Maelstrom 8-Autostage/Maelstrom 9610. B Image of the pulpy dried blood spots (DBS) and iron powder complex isolated by a magnet.

#### Auto-NANDA cost issue

The washed composite was directly subjected to digestion with peptidase (in our case, trypsin). After digestion, the digested peptides released from the composite were separated and retrieved from the iron powder and filter paper debris using the Maelstrom devices. The samples were then desalted simultaneously using a stage-tip adapter for 96 samples. Because peptidase costs approximately 40% of the overall running cost of auto-NANDA, we manually added peptidase using an Eppendorf electronic continuous pipette to reduce the running cost. Although peptidase can be added robotically, it inevitably results in the waste of peptidase in the dead volume of the robotic dispensing system. Therefore, if we add peptidase manually, the cost per sample when simultaneously processing 96 samples is approximately $3, making it acceptable for application in NBS.

#### Comparison and validation of the performances of the auto-NANDA

Next, we compared the performance of SCP, manual NANDA, and auto-NANDA. Manual NANDA identified 3,541 protein groups stably detected in 4 replications, while auto-NANDA identified 4,692 protein groups (Fig 4A). To investigate why auto-NANDA identified more proteins, we compared the total intensities of hemoglobin subunits α (HBA2) and β (HBB), the most abundant proteins in erythrocytes. These proteins can interfere with the detection of low-abundance proteins, as major plasma proteins such as albumin were extensively removed by either method (Fig 4B). In auto-NANDA, the ion intensity of hemoglobins decreased to approximately 60% compared with that in the manual method. The observed signal reduction of hemoglobin likely contributed to the increased detection of low-abundance proteins. In auto-NANDA, the direct collision of the filter paper with the spin tip, along with the periodic reversal of the spin tip’s direction (unique features of Maelstrom 8-Autostage and Maelstrom 9610), likely enhances washing efficiency.

**Figure 4.**
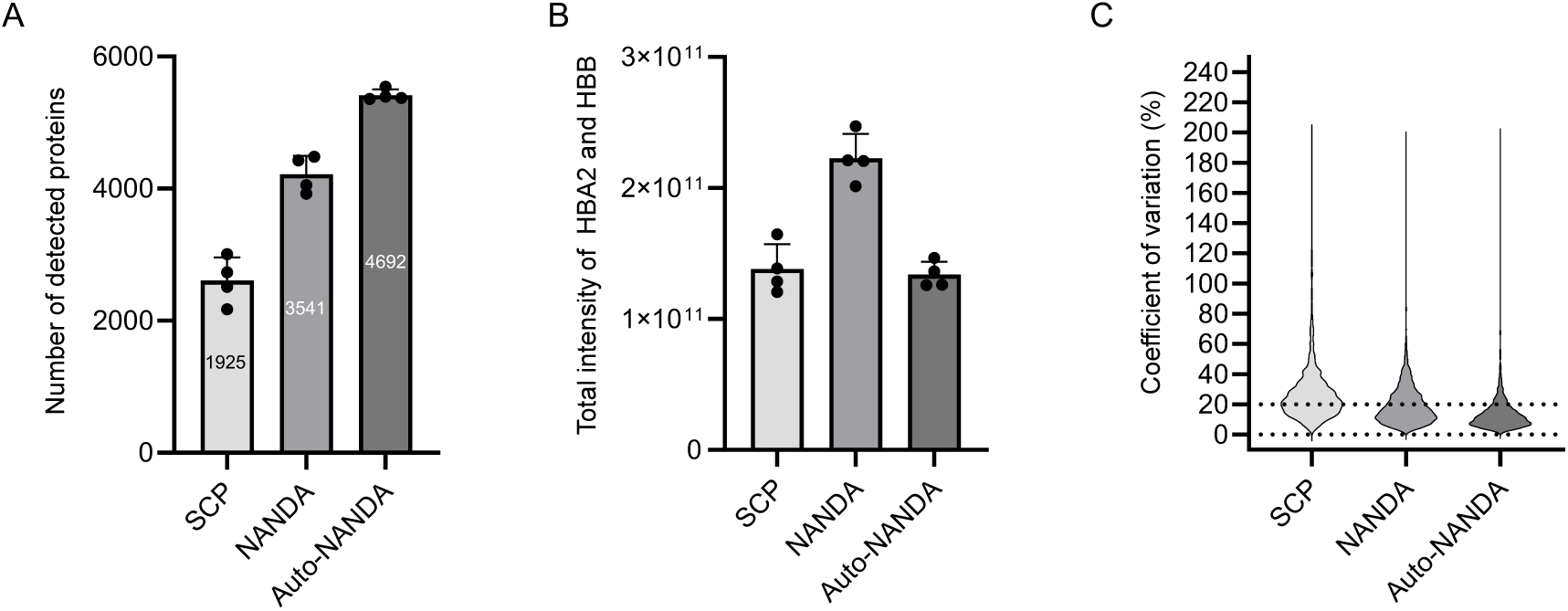
Comparison of proteins detected by manual, non-specifically dried blood spot-absorbed proteins, and automated non-specifically dried blood spot-absorbed proteins. A Bar graph showing the number of proteins detected by sodium carbonate precipitation (SCP), non-specifically dried blood spot-absorbed proteins (NANDA), and automated NANDA (Auto-NANDA) (*n* = 4). The number in the bar graph represents the number of proteins commonly detected in four replicates. B Bar graph showing the total ion intensity of hemoglobin subunits α (HBA2) and β (HBB) detected by sodium carbonate precipitation (SCP), non-specifically dried blood spot-absorbed proteins (NANDA), and automated NANDA (Auto-NANDA) (*n* = 4). C Violin plot showing the distribution of the coefficient of variation (CV) of ion intensities of proteins commonly detected in four replicates.

Additionally, the automated method detected a similar number of proteins from DBS using different filter papers (Whatman 903 Protein Saver Cards; Whatman, Springfield Mill, UK, and PerkinElmer 226 Spot Saver Card; PerkinElmer. Waltham, MA, USA) as it did with the ADVANTEC PKU-S (ADVANTEC, Tokyo, Japan) (Fig EV2). Furthermore, to evaluate the reproducibility of protein detection and quantification between manual and automated methods, the coefficient of variation (CV) values for the ion intensities of individual proteins were analyzed (Fig 4C). The number of proteins with CV values <20% were 2,250 and 3,998 for the manual and automated methods, respectively. Automation facilitated highly reproducible sample preparation, significantly increasing the number of consistently detectable proteins. The lower reproducibility of auto-NANDA than that of conventional proteome analysis using bulk cultured cells (Kawashima *et al*, 2022) may be attributed to sampling variations depending on the punching location of DBS. The results indicated that auto-NANDA considerably improved throughput, proteome depth, and sample preparation robustness, making it suitable for NBS. These benefits ensure consistent and stable results regardless of when, where, or by whom DBS analyses are conducted, which is crucial for clinical testing applications.

### Feasibility of NBS using auto-NANDA

#### Coverage of disease-relevant proteins by auto-NANDA

To assess the applicability of NANDA for NBS, we examined how SCP, manual NANDA, and auto-NANDA detect several disease-relevant proteins in the Online Mendelian Inheritance in Man (OMIM) database (Fig 5). As shown in Fig 5, SCP, manual NANDA, and auto-NANDA detected 975, 1,540, and 1,864 OMIM proteins, respectively, indicating that auto-NANDA enabled us to detect 1.9-fold more OMIM proteins than the SCP method. Because we intend to use this method for NBS in the future, we next compared the detection performance of gene products explicitly listed in the recommended uniform screening panel by the Health Resources & Services Administration (https://www.hrsa.gov/advisory-committees/heritable-disorders/rusp). The data showed that auto-NANDA increased the number of detected gene products to 36, while the SCP method could only detect 17 gene products out of 85 listed gene products (Table EV1). These results implied that auto-NANDA considerably improved the applicability of the non-targeted proteome approach for NBS.

**Figure 5.**
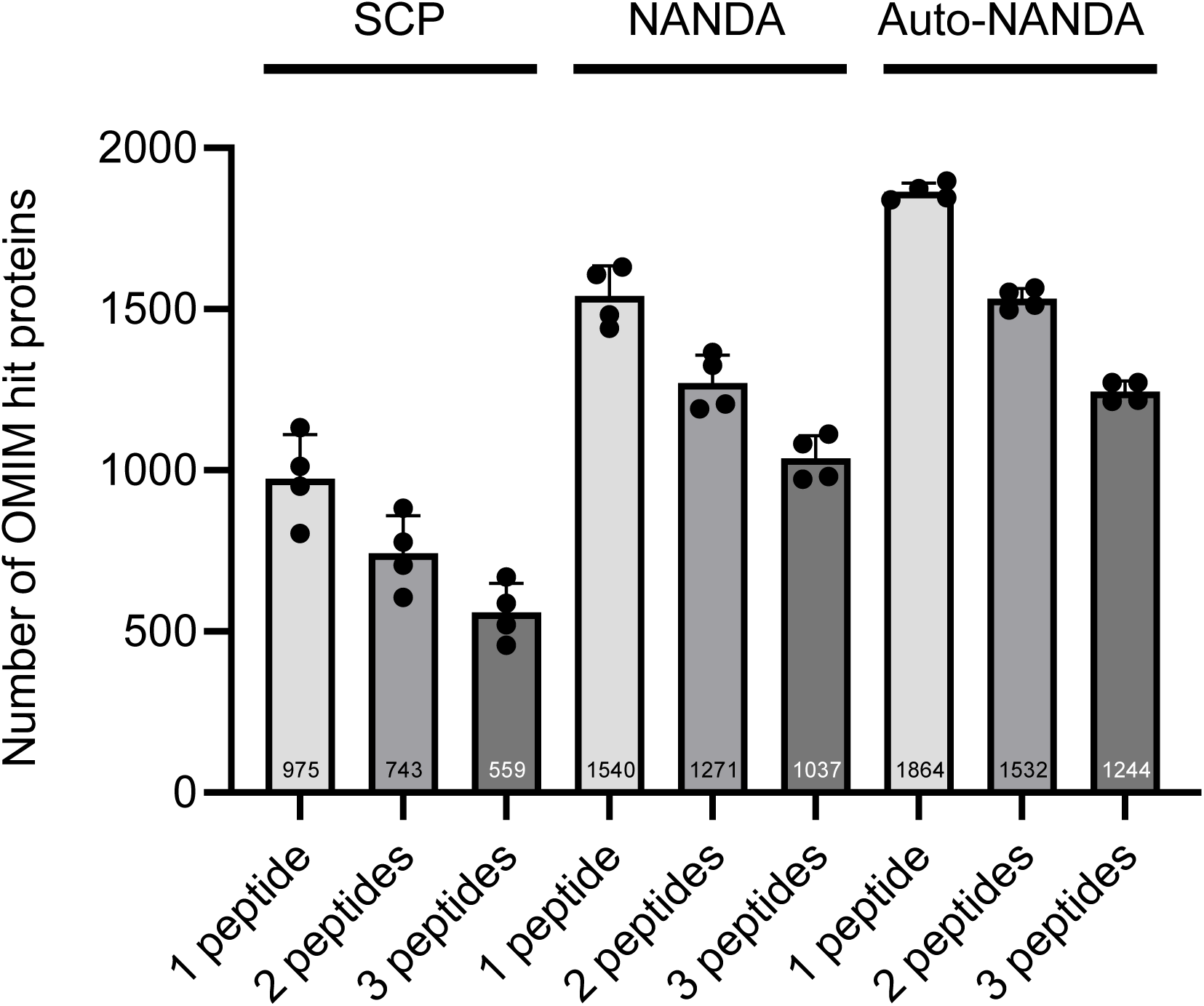
Comparison of the numbers of OMIM hit proteins detected using sodium carbonate precipitation, non-specifically dried blood spot-absorbed proteins (NANDA), and automated NANDA (Auto-NANDA) with peptide number filters from 1 to 3 (*n* = 4). The numbers displayed in the bar graph indicate the average number of proteins detected in each of the four replicate experiments.

In previous discussions of proteins detectable via DBS proteome analysis, we included those identified only by a single peptide. However, for practical NBS applications, single peptide detections are insufficiently robust if they are polymorphic with missense mutations. Therefore, we investigated how the number of proteins detected using two or more peptides varied across SCP, NANDA, and auto-NANDA (Fig 5). The results showed that for the SCP method, 76% (743/975) and 57% (559/975) of proteins were detected using ≥2 and ≥3 peptides, respectively. In manual NANDA, 83% (1,271/1,540) were detected using ≥2 peptides, and 67% (1037/1540) using ≥3 peptides. In Auto-NANDA, 82% (1,532/1,864) were detected using ≥2 peptides and 67% (1244/1864) using ≥3 peptides. Manual and auto-NANDA resulted in a smaller reduction in the number of detected proteins with an increasing number of detected peptides than in the SCP method. Notably, the ATP7B protein, a causative gene product of Wilson disease, was detected only after antibody-assisted enrichment (Collins *et al*, 2021) and only with two peptides using the SCP method. However, it was stably detected with ≥3 peptides using auto-NANDA. In this regard, auto-NANDA detected approximately twice as many OMIM-registered proteins as the SCP method, implying its higher potential for deep screening various genetic disorders (Fig 5). The proteins detected by SCP, manual NANDA, and auto-NANDA are listed in Table EV1.

#### Data reproducibility

Next, we confirmed the daily and inter-device reproducibility of auto-NANDA because reproducibility of the measurement platform is critical for screening (Fig 6). We independently prepared samples from 12 DBS disks using auto-NANDA thrice at different time points (days 1, 32, and 60). These samples were then analyzed using DIA-MS on two different LC-MS instruments with the same specifications and settings. Pearson’s correlation coefficient for the 72 data sets had a median of 0.99 and a minimum of 0.98, demonstrating high reproducibility of overall NANDA (Fig 6). Even if protein extraction was conducted on a different day, Pearson’s correlation coefficient was consistently ≥0.98. This demonstrated the high reproducibility of protein extraction across different days, considering the variability in DBS itself, subsequent protein extraction, protein digestion, desalting, and LC-MS/MS analysis. Additionally, instruments of the same model were confirmed to produce highly correlated data. We thus considered that the overall performance of auto-NANDA coupled with DIA-LC-MS/MS is suitable for large-scale NBS and cohort studies.

**Figure 6.**
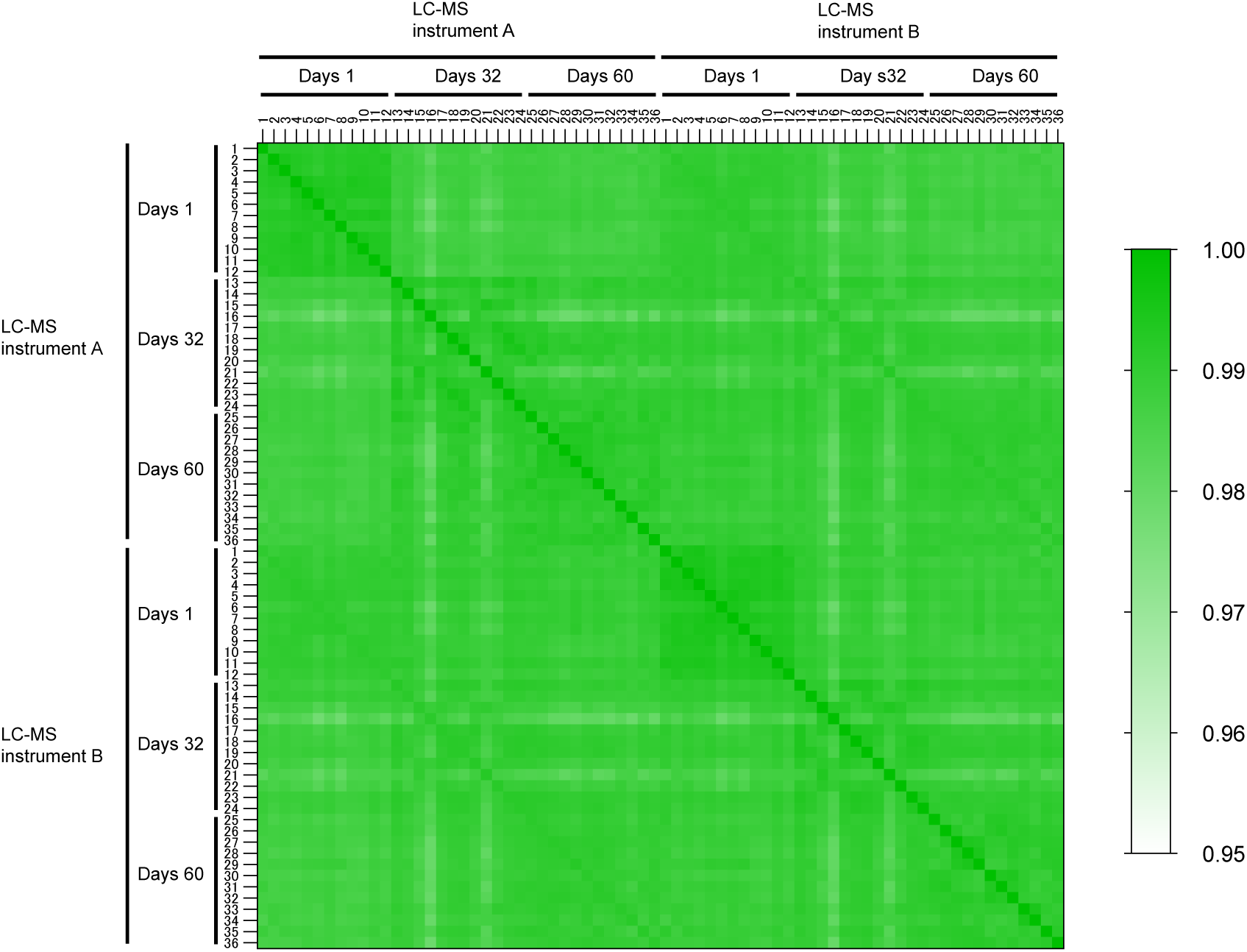
Evaluation of the reproducibility between LC-MS instruments and days in automated non-specifically dried blood spot-absorbed proteins. Pearson’s correlation analysis of protein intensities of samples 1–12, 13–24, and 25–36 prepared using automated non-specifically dried blood spot-absorbed proteins. The method on days 1, 32, and 60 were measured using the same performance LC-MS instruments A and B.

#### Turn-around time of the DIA-LC-MS/MS step

We have already demonstrated the feasibility of NBS using DBS proteome analysis with the SCP method for genetic disorders (McDade *et al*., 2007; Shibata *et al*., 2024). Examining proteins, which are the end products of genes, can complement genetic testing, and the demand for such proteomics approaches is growing. Although the SCP method had throughput- and reproducibility-related issues because of the lack of automation, auto-NANDA solved most of these issues.

One of the remaining issues is the turn-around time of the DIA-LC-MS/MS step. Because non-targeted proteome analysis is conducted one by one, the time required for a single DIA-LC-MS/MS analysis greatly impacts the throughput of NBS. Therefore, we explored the point of compromise between sensitivity and analysis time using the latest Thermo Orbitrap Astral analyzer (Thermo Fisher Scientific, Waltham, MA, USA; Fig 7). Using the Thermo Exploris 480 analyzer (Thermo Fisher Scientific), the number of detected proteins was 5,415 in a 90-min analysis time (12 samples/day, SPD), while the Astral analyzer detected 6,359 proteins in a shorter time of 84 min, i.e., 15 SPD. Additionally, the reduction in the average number of detected proteins when increasing the threshold number of detected peptides to two and three was less pronounced with the Astral analyzer (88% [5,776/6,539] and 76% [4,990/6,539], respectively) than that with the 480 analyzer (78% [4,224/5,415] and 60% [3,265/5,415], respectively). Even when the analysis time was further reduced to 24.5 min (40 SPD), the number of proteins detected with three or more peptides was 3,687, which was higher than the 3,265 proteins detected in the 90-min analysis with the 480 analyzer (Table EV2). These results indicated that the non-targeted proteome analysis of DBS with a protein coverage of >5,000 is feasible at a scale of 10,000 newborns/instrument/year if we use the most advanced instruments.

**Figure 7.**
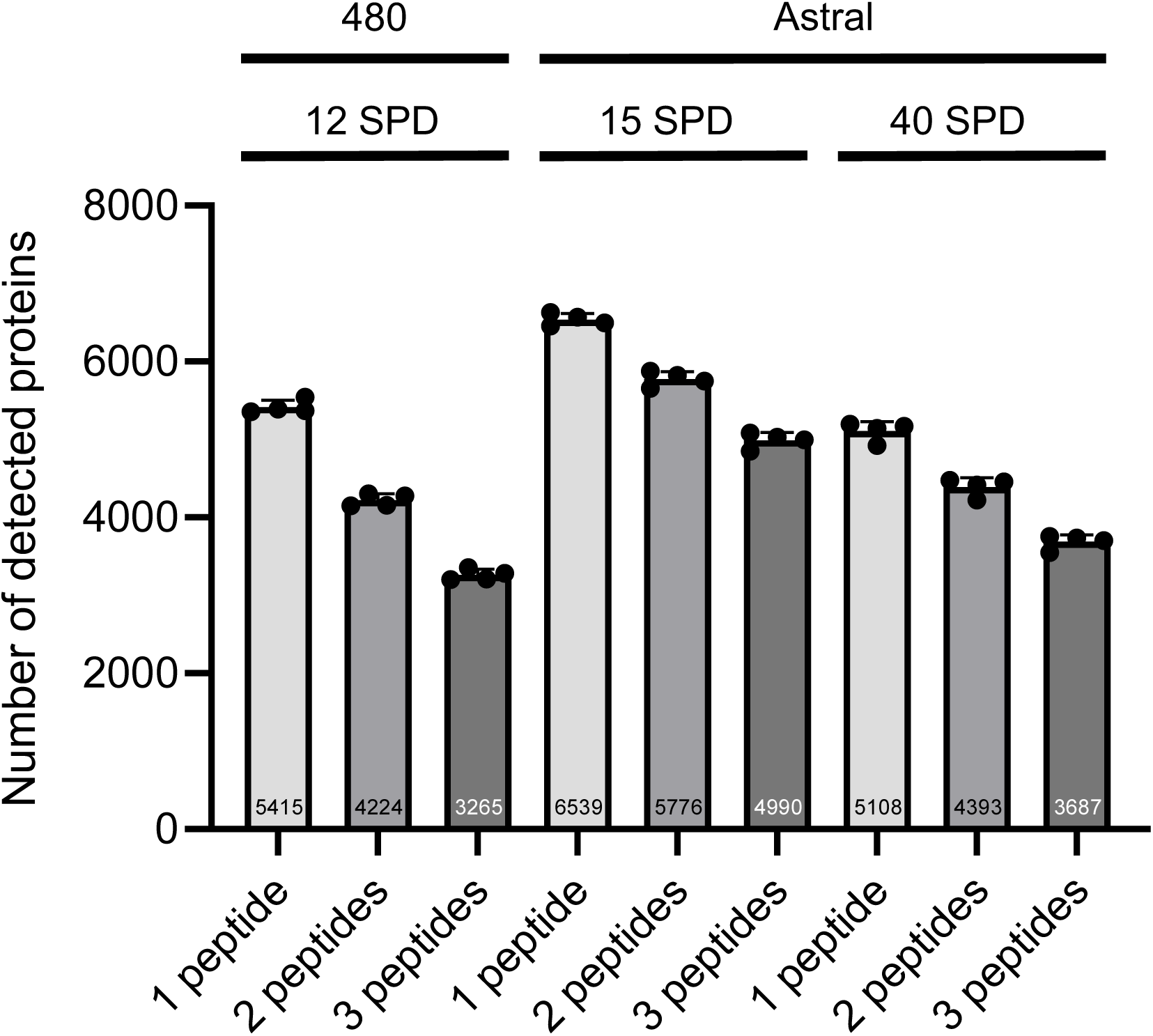
Number of proteins detected using automated non-specifically dried blood spot-absorbed proteins with 480 (12 SPD) and Astral (40 SPD) analyzers and Astral (15 SPD), categorized by the number of detected peptides per protein (*n* = 4). The numbers displayed in the bar graph indicate the average number of proteins detected in each of the four replicate experiments.

#### Current limitations of the non-targeted proteome approach by the DIA-LC-MS/MS

We recently reported the feasibility of NBS using quantitative DBS proteome analysis with the SCP method for genetic disorders (Shibata *et al*., 2024). Although proteome analysis can potentially identify polymorphisms in protein sequences if the protein of interest is sufficiently abundant (Suhre *et al*, 2024), the concept validated by the current study focused solely on quantitative protein profiling. This approach is well-suited for detecting genetic disease caused by mutations resulting in a complete protein-null molecular phenotype. In other words, diseases with dominant inheritance are currently difficult to detect using this approach. Thus, diseases screened using the non-targeted proteome approach should be carefully considered in terms of the mode of inheritance and the abundance of proteins detected. However, despite this limitation, the method provides a fundamental basis for various applications of DBS proteome analysis, including NBS. This is due to its deep protein coverage, which allows it to function as a “one-size-fits-all” platform, reasonable running costs, and fewer ethical concerns than genome sequencing, as long as only quantitative aspects of proteome profiles are analyzed. However, to apply this method in NBS, future studies must thoroughly examine its clinical validity and utility, along with validating its specificity in NBS.

### Conclusions

The non-targeted proteome approach has been demonstrated to possibly contribute to screening patients suffering from inborn errors of immunity with protein-null molecular phenotypes at the neonatal stage (Shibata *et al*., 2024). Building on this study, we developed a robust automated method, auto-NANDA, for pre-processing proteins from DBS to enable NBS for genetic disorders using DBS protein analysis. Using auto-NANDA, the number of detected proteins from DBA increased to >5,000, while our previous SCP method enabled the detection of 2,605 proteins. We demonstrated that auto-NANDA coupled with a sophisticated DIA LC-MS/MS system is cost-effective, highly sensitive, robust, and high-throughput, with minimal peptide detection sensitivity loss. Despite ongoing economic, ethical, and social challenges in implementing proteome-based NBS, auto-NANDA represents a major methodological breakthrough toward realizing this goal.

## Materials and Methods

### Reagents and tools table

#### Ethical considerations for blood sample collection

Written informed consent was obtained from all volunteers prior to participation in the study. The study was conducted in compliance with the Ethical Guidelines for Medical and Biological Research Involving Human Subjects and was approved by the Institutional Ethical Committee of Kazusa DNA Research Institute (Approval No. 2020-06).

### Preparing DBS

DBS were acquired from consenting healthy volunteers via a finger prick onto blood sampling papers (ADVANTEC PKU-S; ADVANTEC; Whatman 903 Protein Saver Cards, Whatman; PerkinElmer 226 Spot Saver Card; PerkinElmer) and dried overnight.

### Protein preparation method (Fig 1)

A 3.2-mm-diameter disk punched out from DBS was added to 500 µL TBST, crushed into pulpy DBS with vigorous agitation for 30 min at maximum using a stirring centrifuge (NSD-12; Nissin Rika Co., Ltd, Tokyo, Japan), and centrifuged for 5 min at 15,000 × *g*. Thereafter, the supernatant was removed, and pulpy DBS was isolated (Step 1,2). Pulpy DBS was washed by adding 500 µL TBST, agitated vigorously for 10 min at maximum using a stirring centrifuge, and centrifuged for 5 min at 15,000 × *g*, after which the supernatant was removed (Step 3). Next, 500 µL 50 mM Tris-HCl pH 8.0 was added, and the mixture was agitated vigorously for 10 min at maximum using a stirring centrifuge and centrifuged for 2 min at 15,000 × *g*, followed by supernatant removal. Afterward, another 500 µL 50 mM Tris-HCl pH 8.0 was added, and the mixture was agitated vigorously using a stirring centrifuge and centrifuged for 5 min at 15,000 × *g*, after which the supernatant was removed to wash the pulpy DBS (Step 4). Subsequently, the pulpy DBS was suspended by adding 200 µL 50 mM Tris-HCl pH 8.0 and agitating vigorously for 3 min at maximum using a stirring centrifuge (Step 5).

### Automated protein preparation using NANDA (Fig 3 and Video EV)

Ninety-six well plates for Maelstrom 8-Autostage/Maelstrom 9610 instrument (TANBead) processing were prepared. Briefly, 500 µL 1× TBST, 5 mg freshly prepared iron powder solution (3–5 µm particle size, 5 mg/25 µL in 50% glycerol), and one 3.2- mm diameter disk punched out from DBS were added to the first row/first 96-well plate. Next, 500 µL 1× TBST was added to the second row/second 96-well plate, and 500 µL 50 mM Tris-HCl pH 8.0 was added to the third and fourth row/third and fourth 96-well plate. Finally, 200 µL of 50 mM Tris-HCl pH 8.0 was added to the fifth row/fifth 96-well plate. Subsequently, the 96-well plate was set on the Maelstrom 8-Autostage/Maelstrom 9610, and the protein extraction procedure from DBS was initiated. A spin tip was inserted into the first row/plate, and the DBS was crushed by agitating in TBST for 30 min at 3,500 rpm with inverting every 10 s. This formed a complex of pulp-like DBS and iron powder. Next, a magnetic rod was inserted into the spin tip, and the complex was adsorbed and collected on the spin tip over 60 s before being moved to the second row/second plate containing TBST. After removing the magnet rod from the spin tip in TBST, the complex was washed by agitation for 10 min at 3,500 rpm with inverting every 10 s. Afterward, the complex was transferred to the third row/third plate containing 50 mM Tris-HCl pH 8.0 and washed by agitating the spin tip for 10 min at 3,500 rpm with inverting every 10 s. This washing process was repeated in the fourth row/fourth plate containing 50 mM Tris-HCl pH 8.0 by agitating the spin tip for 2 min at 3,500 rpm with inverting every 10 s. Finally, the complexes were transferred to the fifth row/fifth plate containing 50 mM Tris-HCl pH 8.0 and suspended by agitating the spin tip for 2 min at 3,500 rpm with inverting every 10 s.

### Protein digestion

For protein digestion, 1 µg trypsin/Lys-C Mix (CAT# V5072; Promega, Madison, WI, USA) was gently mixed with the sample for 14 h at 37 °C. The digested sample was then acidified with 50 µL 5% trifluoroacetic acid (TFA) and sonicated using a Bioruptor II (CosmoBio, Tokyo, Japan) at a high level for 5 min at 24 °C. Thereafter, the sample was desalted using a styrene-divinylbenzene polymer (SDB) stop-and-go extraction (STAGE) tip (SDB-STAGE tip; GL Sciences, Tokyo, Japan), which was washed with 25 µL 80% acetonitrile (ACN) in 0.1% TFA, followed by equilibration with 50 µL 3% ACN in 0.1% TFA. Next, the sample was loaded onto the tip, washed with 80 µL 3% ACN in 0.1% TFA, and eluted with 50 µL 36% ACN in 0.1% TFA. The eluate was dried in a centrifugal evaporator (miVac Duo concentrator; Genevac, Ipswich, UK). The dried sample was then redissolved in 0.02% decyl maltose neopentyl glycol (Anatrace Products, LLC, Maumee, OH, USA). Subsequently, the redissolved sample was assayed for peptide concentration using a Lunatic instrument (Unchained Labs, Pleasanton, CA, USA) and transferred to an LC vial (Thermo Fisher Scientific). Additionally, we omitted the reduction and alkylation steps to streamline the process. Although the number of identified peptides slightly decreased without these steps, the number of identified proteins did not decrease (Fig EV3). Considering the need to process many samples, skipping the reduction and alkylation steps was beneficial.

### LC-MS/MS with DIA

The redissolved peptides were injected directly onto a 75 µm × 25 cm nanoLC column (Aurora C18, particle size 1.6 µm, 120 Å; IonOpticks, Fitzroy, VIC, Australia) at 60 °C and separated using a 90-min gradient [A = 0.1% formic acid (FA) in water, B = 0.1% FA in 80% can] with the following settings: 0 min at 4% B, 78 min at 32% B, 84 min at 75% B, and 90 min at 75% B, all at a flow rate of 200 nL/min. This separation was performed using an UltiMate 3000 RSLCnano LC system (Thermo Fisher Scientific). The eluting peptides from the column were then analyzed on an Orbitrap Exploris 480 equipped with an InSpIon system (Thermo Fisher Scientific) (Kawashima *et al*, 2023). For DIA, the precursor range was defined based on previously established parameters (Kawashima *et al*., 2022). MS1 spectra were collected at nine different ranges corresponding to the DIA method (395–605, 395–705, 395–805, 495–705, 495–805, 495–905, 595–805, 595–905, 695–905 m/z) at a resolution of 15,000, with an automatic gain control target of 3 × 10^6^ and a maximum injection time of “Auto.” MS2 spectra were collected in the range of 200–1,800 m/z at a resolution of 30,000, with an automatic gain control target of 3 × 10^6^, a maximum injection time set to “Auto,” and a stepped normalized collision energy of 22, 26, and 30%. For the DIA window patterns (400–600, 400–700, 400–800, 500–700, 500–800, 500–900, 600–800, 600–900, 700–900 m/z), an optimized window arrangement was employed using Xcalibur 4.3 (Thermo Fisher Scientific). The isolation width for MS2 was set to 4, 6, and 8 Th at 200, 300, and 400 m/z with the DIA mass range, respectively.

In the analysis using Orbitrap Astral (Thermo Fisher Scientific), the redissolved peptides were injected directly onto a 75 µm × 30 cm nanoLC column (ReproSil-Pur C18, particle size 1.5 µm, 100 Å; CoAnn Technologies, Richland, WA, USA) at 60 °C, and separated using a 84.5-min gradient (15 SPD) comprising 1% B at a flow rate of 600 μL/min in 0–0.5 min, 1–8% B at a flow rate of 600–200 μL/min in 0.5–4.5 min, 8–23% B at a flow rate of 200 μL/min in 4.5–60 min, 23–40% B at a flow rate of 200 μL/min in 60–78.5 min, 40–98% B at a flow rate of 200 μL/min in 78.5–79.5 min, 98% B at a flow rate of 200 μL/min in 79.5–80.5 min, 98% B at a flow rate of 200-600 μL/min in 80.5–81.5 min, 98% B at a flow rate of 600–750 μL/min in 81.5–82.5 min, and 98% B at a flow rate of 750 μL/min in 82.5–84.5 min, and a 24.5-min gradient (40 SPD) comprising 1% B at a flow rate of 600 μL/min in 0–0.5 min, 1–11% B at a flow rate of 600–300 μL/min in 0.5–3.5 min, 11–23% B at a flow rate of 300 μL/min in 3.5–15.5 min, 23–40% B at a flow rate of 300 μL/min in 15.5–19.5 min, 40–98% B at a flow rate of 300 μL/min in 19.5–20.5 min, 98% B at a flow rate of 300 μL/min in 20.5–21.5 min, 98% B at a flow rate of 300–600 μL/min in 21.5–22.5 min, 98% B at a flow rate of 600–750 μL/min in 22.5–23 min, and 98% B at a flow rate of 750 μL/min in 23–24.5 min. Subsequently, the eluting peptides from the column were analyzed on an Orbitrap Astral (Thermo Fisher Scientific) equipped with an InSpIon system. MS1 spectra were collected in the range of m/z 380–980 at a 240,000 resolution using Orbitrap to set an automatic gain control target of 500% and a maximum injection time of 5 ms. MS2 spectra were collected at m/z 250– 2,000 using an Astral analyzer (Thermo Fisher Scientific) to set an automatic gain control target of 600 and 500%, maximum injection time of 4 and 3 ms for 15 and 40 SPD, respectively, and a normalized collision energy of 25%. The isolation width for MS2 was set to 1.6 and 2 Th for 15 and 40 SPD, respectively, with window placement optimization turned on.

### Data analysis

The DIA-MS data were queried against the *in silico* human spectral library using DIA-NN v.1.9.1 (https://github.com/vdemichev/DiaNN) (Demichev *et al*, 2020). Initially, the spectral library was generated from the human protein sequence UniProt database (proteome ID UP000005640, 20,598 entries, downloaded on April 1, 2024) using DIA-NN. The parameters for generating the spectral library were as follows: digestion enzyme, trypsin; missed cleavages, 1; peptide length range, 7–45; precursor charge range, 2–4; fragment ion m/z range, 200–1,800. The precursor mass range was varied according to the DIA method (m/z 400–600, 400–700, 400–800, 500–700, 500–800, 500–900, 600–800, 600–900, 700–900). Additionally, “FASTA digest for library-free search/library generation,” “deep learning-based spectra, RTs and IM prediction,” and “n-term M excision” were enabled. For the DIA-NN search, the following parameters were applied: a mass accuracy of 10 ppm, MS1 accuracy of 10 ppm, protein inference based on genes, utilization of neural network classifiers in single-pass mode, quantification strategy using QuantUMS (high precision), and cross-run normalization set to “RT-dependent.” Furthermore, “Unrelated runs,” “Peptideforms,” “Heuristic protein inference,” and “No shared spectra” were enabled, whereas the use of match-between-run was deactivated. The threshold for protein identification was set to 1% or less for both precursor and protein false discovery rates. Protein quantification values were aggregated over the quantification values of unique peptides as calculated by DIA-NN.

Log2 transformation of protein intensity was applied, and a filtering step was conducted to ensure that for each protein, at least one group contained a minimum of 70% valid values. Missing values were imputed using random numbers drawn from a normal distribution (width, 0.3; downshift, 1.8) using Perseus v1.6.15.0 (Tyanova *et al*, 2016), and Pearson’s correlation analysis was performed. Gene Ontology enrichment analysis for membrane proteins and matching to the OMIM database were conducted using DAVID (https://david.ncifcrf.gov/tools.jsp). The analysis of the origin of blood components of proteins was based on our previous report (Nakajima *et al*., 2020). Other graphs were generated using GraphPad Prism 9 (GraphPad Software, San Diego, CA, USA) or Excel (Microsoft Corporation, Redmond, WA, USA).

## Supporting information

https://docs.google.com/spreadsheets/d/11os2roULB0BS5rw1ZKP2WCXN1OTAgupt/edit?usp=sharing&ouid=103067625253435575273&rtpof=true&sd=true

https://docs.google.com/spreadsheets/d/1f071lcjqjT0BGO2rd2ucFHjCoXiV3SZB/edit?usp=sharing&ouid=103067625253435575273&rtpof=true&sd=true

https://docs.google.com/spreadsheets/d/1-cerYQqCHtrQxKvQa7I52nR009-aaKrx/edit?usp=sharing&ouid=103067625253435575273&rtpof=true&sd=true

## Acknowledgments

This study was supported in part by JSPS KAKENHI under Grant Numbers 21K07877 and 24K11013 and by AMED under Grant Number JP22ek0109586. This work was also supported by the Kazusa DNA Research Institute Foundation.

## Author Contributions

D.N. and Y.K. designed the study; D.N. performed research; M.I. and R.K. analyzed data; D.N. and H.S. provided reagents and samples; D.N., O.O., and YK wrote the paper with input from all authors.

## Data Availability

The LC-MS/MS data are available from the ProteomeXchange Consortium via the jPOST partner repository under the accession codes PXD058804.

---

**Figure EV1.**
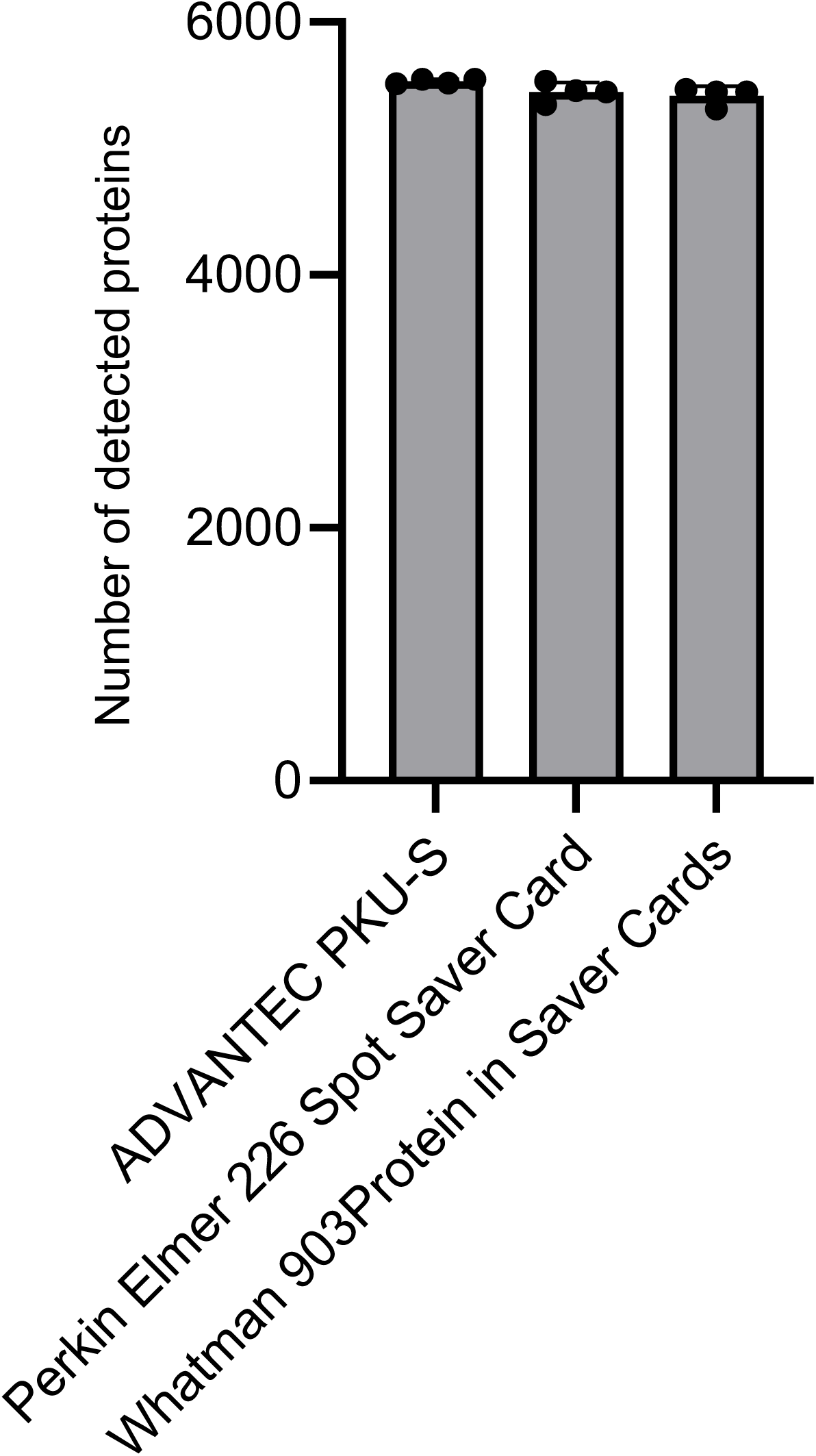
Comparison of the number of proteins detected from automated non-specifically dried blood spot-absorbed proteins using filter paper manufactured using three makers (*n* = 4).

**Figure EV2.**
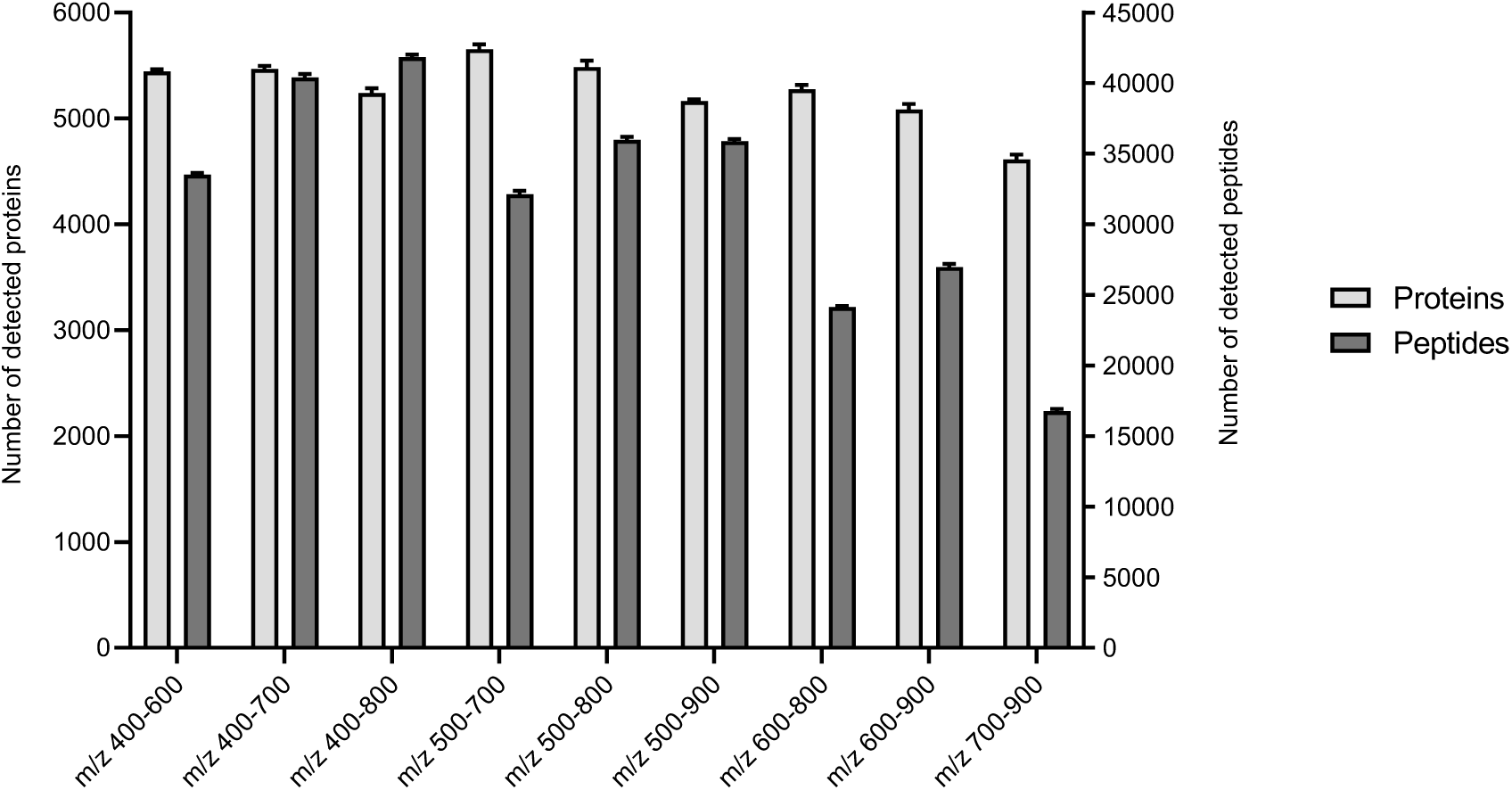
DIA-MS m/z range optimization for the proteome analysis of automated non-specifically dried blood spot-absorbed proteins. Bar graph showing the number of proteins and peptides detected in each m/z range. The left and right *y*-axes indicate the number of proteins and peptides, respectively (*n* = 4).

**Figure EV3.**
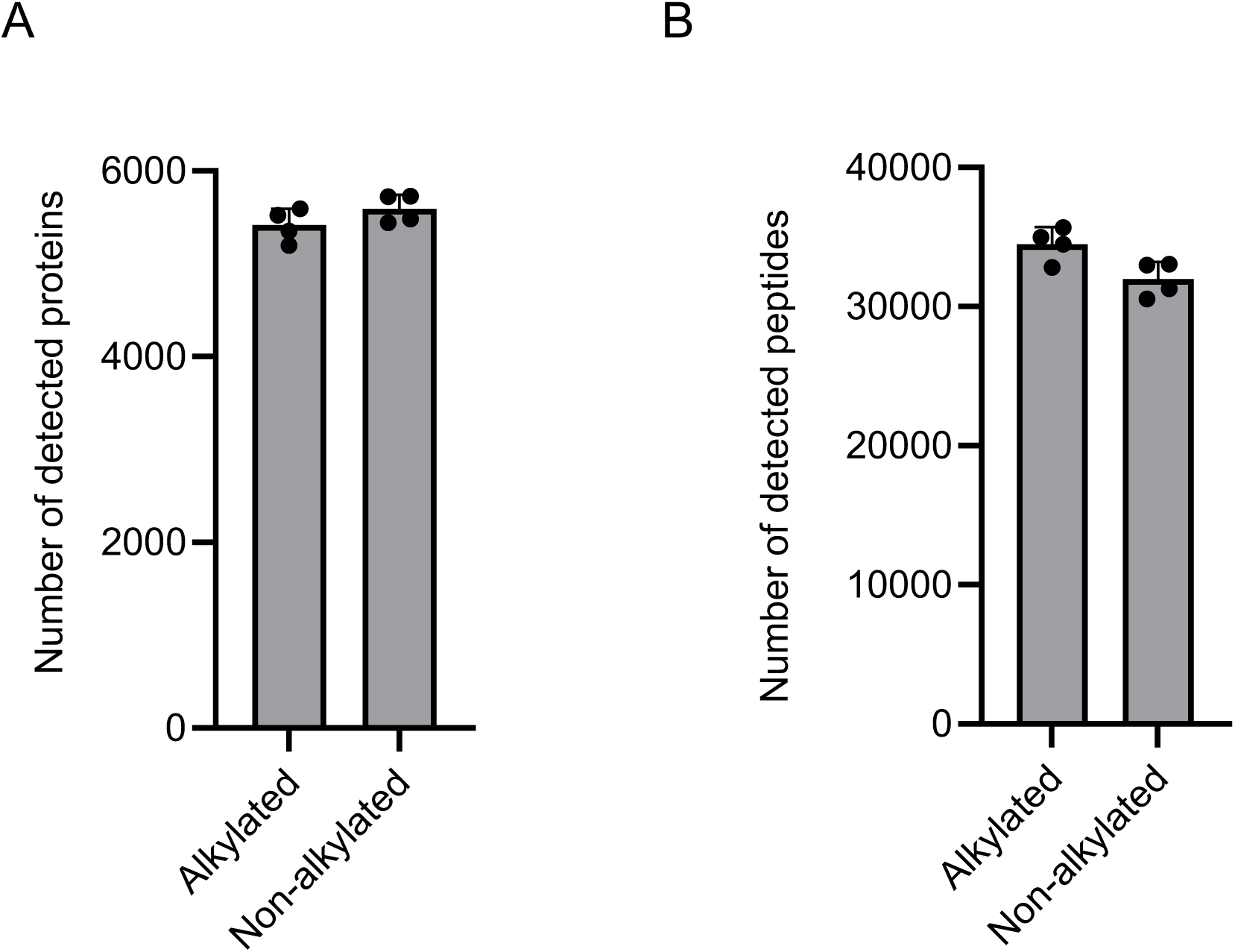
Comparison of the number of proteins and peptides detected using automated non-specifically dried blood spot-absorbed proteins with or without alkylation treatment. A Bar graph showing the number of proteins detected with or without alkylation treatment of automated non-specifically dried blood spot-absorbed proteins (NANDA) (*n* = 4). Alkylation was performed after trypsin digestion of NANDA extracted using an automated extraction method. 2-chloroacetamide and tris (2-carboxyethyl) phosphine were added to final concentrations of 40 and 10 mM, respectively, and the mixture was incubated at 80 °C for 15 min. B Bar graph showing the number of peptides detected with or without alkylation treatment of auto-NANDA (*n* = 4).

